# Anti-CRISPR RNAs: designing universal riboregulators with deep learning of Csy4-mediated RNA processing

**DOI:** 10.1101/2020.11.15.384107

**Authors:** Haotian Guo, Xiaohu Song, Ariel B. Lindner

## Abstract

RNA-based regulation offers a promising alternative of protein-based transcriptional networks. However, designing synthetic riboregulators with desirable functionalities using arbitrary sequences remains challenging, due in part to insufficient exploration of RNA sequence-to-function landscapes. Here we report that CRISPR-Csy4 mediates a nearly all-or-none processing of precursor CRISPR RNAs (pre-crRNAs), by profiling Csy4 binding sites flanked by > 1 million random sequences. This represents an ideal sequence-to-function space for universal riboregulator designs. Lacking discernible sequence-structural commonality among processable pre-crRNAs, we trained a neural network for accurate classification (f1-score ≈ 0.93). Inspired by exhaustive probing of palindromic flanking sequences, we designed anti-CRISPR RNAs (acrRNAs) that suppress processing of pre-crRNAs via stem stacking. We validated machine-learning-guided designs with >30 functional pairs of acrRNAs and pre-crRNAs to achieve switch-like properties. This opens a wide range of plug-and-play applications tailored through pre-crRNA designs, and represents a programmable alternative to protein-based anti-CRISPRs.

## Main text

Benefiting from the simplicity of base pairing, RNA-based devices can be designed *de novo* to enable post-transcriptional regulations and thus received much attention in past decades^1–4^. Though protein-based transcriptional factors have been commonly adopted in genetic circuits, the limited number of available components hinders the applicability of more complex systems^5^. As an alternative, riboregulators consisting of switch and trigger RNAs^1, 2^, can be theoretically designed within an enormous sequence space. They may afford large sets of orthogonal components^5^, where switches only interact with cognate triggers and not the others in the set.

To this end, numerous artificial riboregulator designs have been developed, such as antisense RNAs^6, 7^, cis-repression^8^, toehold switches^9^, and looped antisense oligonucleotides (LASO)^10^ (Fig. S1a-d). We refer here to switch RNAs domains that sense trigger RNAs with arbitrary sequences, and that drive output signals with invariable sequences, as sensors and effectors, respectively. Earlier riboregulators^8, 11^ were derived from natural systems having overlapping or base-pairing sensor and effector domains, leading to significant design constraints (Fig. S1a, b). More recent translational riboregulators reduced such constraints by separating sensors and effectors^9, 10, 12^ (Fig. S1c, d). However, there are still obstacles to generalizing the utility, hitherto limited to relatively few reusable, high-performance parts. Effectors engineered from native ribosomal binding sites (RBS)^6–10, 12^ or transcriptional terminators^11^ are not adaptable for different mechanisms or across organisms. Without overlaps with sensors, the effectors were regulated by flanking structural changes (Fig. S1c, d), yet the nature of which was not fully understood in advance of riboregulator designs. As a result, different designs displayed large variations of performances, with disparate open ranges of ON and OFF states (Fig. S1e)^9, 10, 12, 13^.

To overcome these hurdles, here we propose a roadmap for a universal design of riboregulators from first principles (Fig. 1). As its core, the effector domain is functionalized by factors orthogonal from native physiologies, and its outputs flexibly adapted into various downstream applications. Next, it should be separated from sensor domains. Switch-trigger interactions do not involve the effector. Rather, triggers are “anti-flanking” RNAs that only bind to the flanking context of the effector. Therefore, the effector needs to be sensitive to such context changes. Moreover, its outputs should span in a closed interval, and ideally be binary, so that different variants within a particular riboregulator family could perform within comparable ranges of maximal and minimal outputs. We refer to having such unprecedented binary outputs in response to changing flanking context, as “context-ultrasensitivity”.

**Fig. 1.**
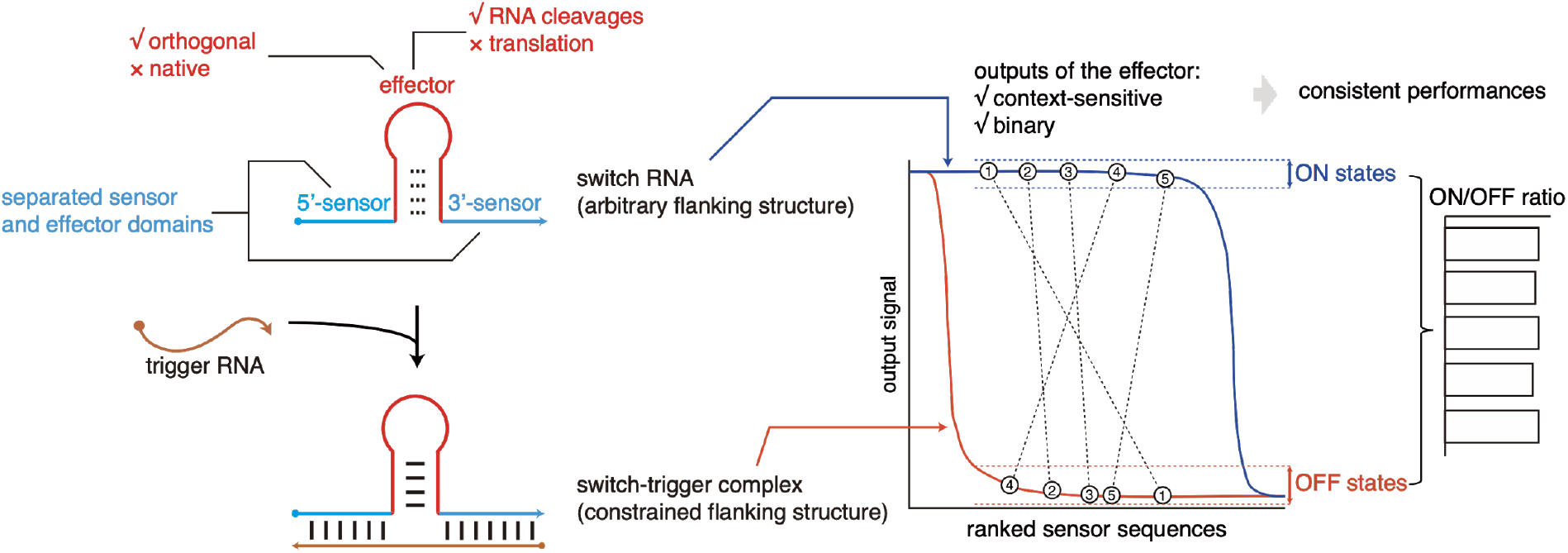
A universal design of riboregulator. Switch RNAs of riboregulators consist of two domains, sequence-variable sensor domain and invariable effector domain. The function of the effector should be realized by heterogeneous factors that are orthogonal from native physiologies such as translation. The output of the effector domain needs to be adaptable for downstream events, *e.g.,* RNA-level signal or RNA cleavages. To minimize constraints of usable sensor sequences, the effector (in red) should be decoupled from the sensor domains (in blue), and should not be required to fold into a specific structure (implied by dashed lines). As triggers (in brown) are anti-flanking RNAs that only aim to convert the flanking context of the effector from random structures into a specific one (base pairing in solid lines), the effector needs to be highly sequence- or structural-sensitive to functionalize such process. Illustrated on the right, the ideal outputs should be mostly binary in response to sequence and structure flanking contexts, to enable consistent performances across riboregulators with different sensor sequences (shown in 5 hypothetical variants).

To apply these principles, we hypothesized that the CRISPR/Csy4(Cas6f) mediated RNA processing^14^ would be an ideal candidate. In *Pseudomonas aeruginosa*, the single-turnover endoribonuclease Csy4 cleaves CRISPR RNA precursors (pre-crRNAs) into mature CRISPR RNAs (crRNAs) for bacterial immunity against foreign genetic elements. Csy4-mediated processing has been routinely used in heterogeneous organisms^15–18^ to create RNA cleavage that is highly adaptable for downstream regulations^15, 16, 19–21^. Crystal structures of Csy4-crRNA complexes reveal a “cleavage fork” geometry where the 5’ and 3’ ends of the Csy4 binding site (CBS) are separated by the Csy4’s RNA-binding domain^22^ (Fig. S2a). This implies that the cleavage requires specific flanking structures. This is in line with *in vitro* results where flanking CBS by two additional G–C pairs led to ∼1500-fold slower processing of pre-folded RNAs^23^. Similar geometries were also commonly reported among other Cas6-crRNA homologs, suggestive of conservative context sensitivity^24^. Finally, the percentage of processed RNAs could only vary from 0 to 100%, i.e. within a closed interval.

Here we characterize Csy4-mediated processing in *Escherichia coli* to derive design principles of its riboregulators. Using massively parallel reporter assays^25–32^, we show a nearly “all-or-none” activity dependency on the flanking context of CBS, without trivial sequence or structural motifs, therefore satisfying the aspired novel context-ultrasensitivity. By brute-force testing of palindromic flanking sequences, we confirm that 6-base pairing (bp) helices sufficiently inhibit pre-crRNAs processing. Accordingly, we propose a class of anti-CRISPR RNAs (acrRNAs) that repress processing of cognate pre-crRNA variants by forming stacked junctions with the flanking strands of CBS. To identify processable and thus switchable pre-crRNAs, we trained a 2-dimensional convolutional neural network Seq2DFunc^33^ with high accuracy. We demonstrate Seq2DFunc-aided design pipeline by validating over 30 designs in *E. coli*. We show functional systems with different sequences can similarly control gene expression from negligible to full induction, suggesting high performance and consistency. Thus, we anticipate that acrRNAs can be a potential solution for universal riboregulators, as well as an efficient alternative of CRISPR-Cas protein inhibitors (*i.e.,* anti-CRISPR or acr proteins)^34, 35^ to control CRISPR-Cas systems. In addition, we expect that our exploration of generating a unified pipeline to design riboregulators for any sequences, would give novel insights of computational designs in synthetic biology, and our study on CRISPR processing mechanisms may inspire new CRISPR-driven methodologies.

## Results

### Coding sequence cleavage assay to report Csy4’s activity in vivo

A gene silencing reporter system was previously designed to follow Csy4-mediated cleavage of CBS in *E. coli* by inserting the substrate of interest in-frame after the gene’s start codon^15^. In the original assay, sequence changes near RBS could contribute to variations of reporter gene expression^36^. Thus, we inserted the substrate of interest in-frame within a GGGS linker between sequences encoding AraC and green fluorescent protein (GFP) (Fig. 2a). Hereafter, RNAs containing the substrate are termed pre-crRNA in general. We validated the new coding sequence cleavage assay with pre-crRNA containing 28-nt wild-type CRISPR repeat (pre-crRNA-wt), where overexpression of Csy4 specifically reduced GFP production to autofluorescence levels (Fig. 2b,c).

**Fig. 2.**
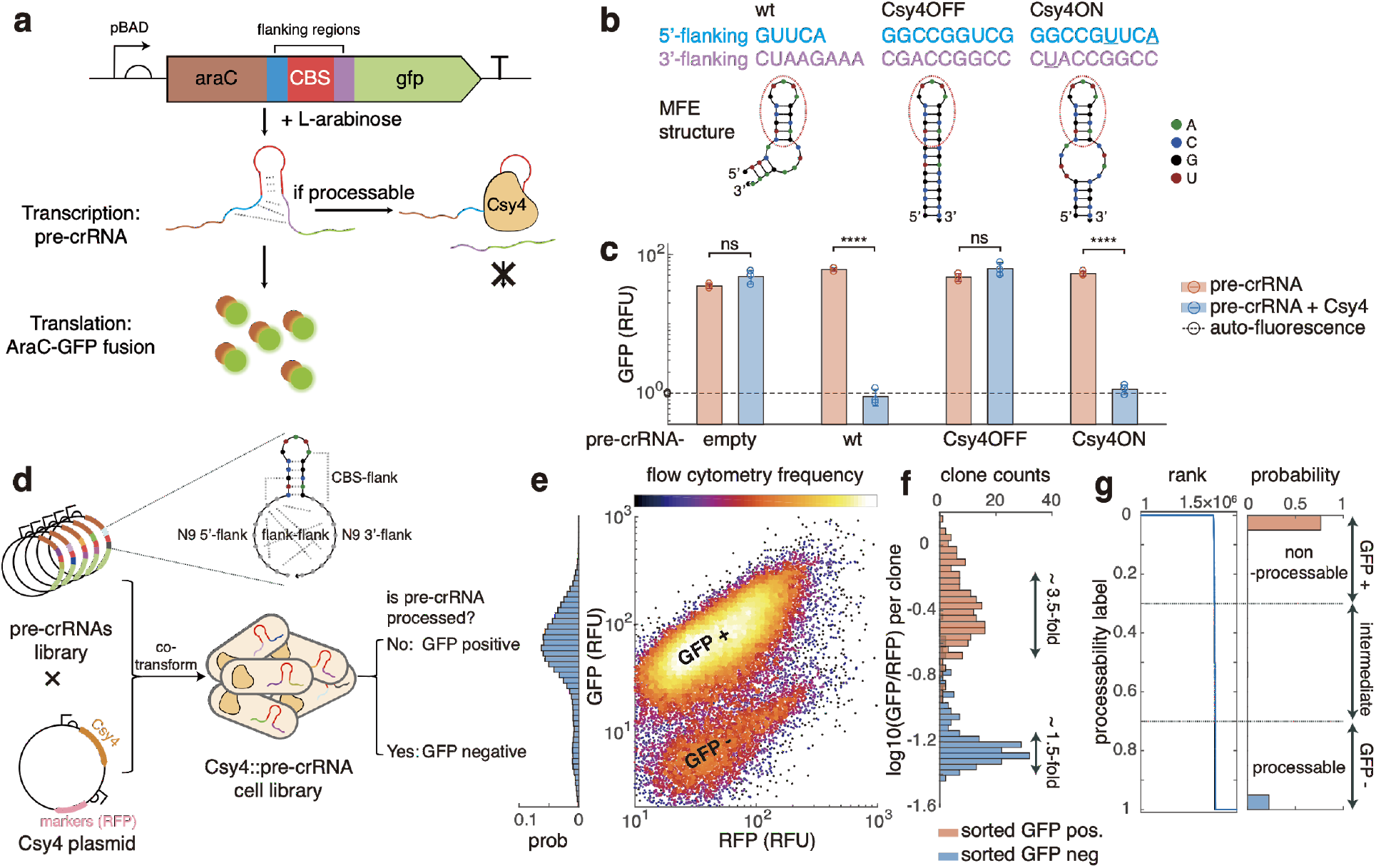
Context-ultrasensitivity of Csy4-mediated RNA processing. **a**, Coding sequence cleavage assay to characterize Csy4-mediated processing CBS with random or designed flanking sequences is inserted between sequences encoding AraC and GFP. If Csy4 processes the inserted CBS, the mRNA is cleaved and thereby cannot be translated, resulting in reduced GFP production. **b**, Sequence and predicted secondary structures of wild-type, Csy4OFF and Csy4ON substrates. **c**, Processing of wild-type, Csy4OFF and Csy4ON substrates, measured by GFP levels without (red) or with (blue) Csy4 expression. RFU, relative fluorescence unit, the ratio of arbitrary units between measured sample and autofluorescence. Error bars, standard deviation of three biological replicates. ****, p-value < 0.0001, and ns, not significant (p-value > 0.05), measured by paired Student’s t-test. **d**, Characterization of random-flanking pre-crRNA libraries. CBS flanked by random sequences were inserted within araC-GFP sequence. Resulted pre-crRNA libraries were co-transformed with Csy4 plasmids constitutively expressing Csy4, and RFP for selection. Cell libraries were sorted by fluorescence to identify processable and non-processable pre-crRNAs. **e**, GFP versus RFP expression levels of cells coexpressing Csy4 plasmids, and pre-crRNA library 1, showing two distinct subpopulations. **f**, GFP over RFP ratios of isolated single colonies sorted from GFP positive and negative subpopulations. Coefficients of variations among two subpopulations were measured as 3.5- and 1.5-fold, respectively. **g**, Processability labeled by next-generation sequencing results. Shown from left to right are ranked labels and distribution of labels. Thresholds for being processable and non-processable are arbitrarily defined as > 0.7 and < 0.3.

### Csy4 is unusually ultrasensitive to the flanking structures of CBS

To probe context sensitivity in vivo, we characterized rationally-designed pre-crRNA variants with our cleavage assay. Intuitively, we speculated that CBS flanked by two long-enough complementary strands cannot be accessed by Csy4. We therefore designed a substrate termed Csy4OFF to flank CBS by 9-bp stem, and a substrate termed Csy4ON with a relaxed structure by flanking CBS with 4-nt loops on each side and then a 5-bp proximal stem (Fig. 2b), as predicted by Nupack^37^. As expected, Csy4OFF and Csy4ON presented repressed and wild-type level processing, respectively (Fig. 2c). We extended the characterization to 8 different rationally designed variants, yet many of the relaxed structures did not recover cleavage (Fig. S2b, and Table S1). Thus, Csy4 is highly selective to the flanking structure context of CBS in vivo.

We then estimated Csy4’s selectivity to arbitrary flanking contexts, using pre-crRNA synthesized libraries, containing 9 random nucleotides on each side of the CBS (i.e. N9-CBS-N9), within the gene silencing reporter system (Fig. 2d). Base pairings between flanking regions or flank-to-CBS were potentially allowed. We observed bimodal distribution of GFP levels when co-expressing the pre-crRNA libraries with Csy4 (Fig. 2e and S3a) in experiments from independently cloned libraries, covering about 3 million individuals in total (Fig. S3b). Using fluorescence activated cell sorting (FACS) followed by isolation of 192 clones from each subpopulation, we validated that Csy4 indeed responds to flanking context in an “all-or-none” binary manner (Fig. 2f), where variation among GFP-positive clones can be explained by changing stabilities of pre-crRNAs. Then, by next-generation sequencing of sorted clones (i.e. FACS-seq^25–27, 30, 32^), we identified about 1 million distinct pre-crRNAs. In line with the flow cytometry observation, most sequences were dominantly detected either in the GFP positive or negative subpopulation (Fig. S3c), thus further labeled as processable or non-processable pre-crRNAs (Fig. 1g, and S3d). Previously characterized genetic elements usually exhibit continuous landscapes of flanking sequences to functions^30, 36, 38^, for instance, the thermodynamic relationships between the flanking contexts of Shine–Dalgarno sequence and RBS strengths^39^. This is in contrast to the sequence context-ultrasensitivity of Csy4-mediated processing, which makes CBS a better effector domain.

### Sequence-independent repression of processing by adjacent stems

Next, we sought mechanisms to switch processability by trigger-switch interactions. We asked whether our intuition regarding inhibiting cleavage by flanking CBS with complementary strands is generally true. For the sake of simplicity, we only considered adjacent stems without bulges or internal loops. Among profiled pre-crRNAs, longer adjacent stems indeed indicated higher possibility of inhibition (Fig. 3a).

**Fig. 3.**
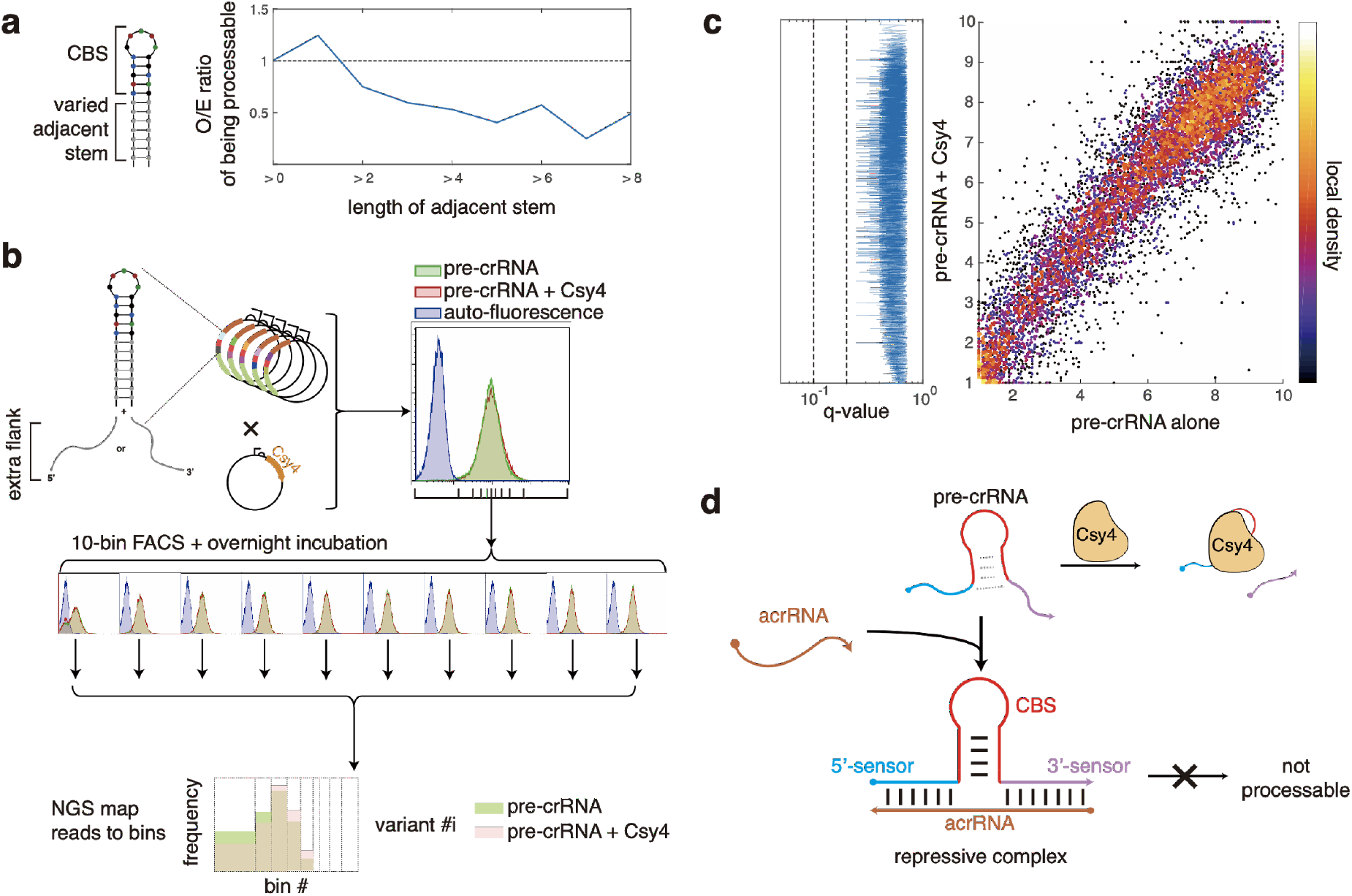
Sequence-independent repression by stem stacking. **a**, Longer adjacent stems reduced the probability of being processed in profiled random-flanking variants. O/E ratio, the ratio of observed probability of being processed under the condition that the adjacent stem is longer than the base pairs on x-axis, over expected probability of being processed among all profiled variants. **b**, Characterization of synthesized palindromic-flanking library. CBS flanked by 6- or 9-bp palindromes followed with an extra 5’- or 3’-unstructured linker to avoid the bias from vector, were inserted within araC-GFP, and co-transformed with Csy4 plasmid for characterization. With (red) or without (green) Csy4, the cell library displayed indistinguishable GFP expression distributions, significantly higher than autofluorescence (blue). Cell libraries were further sorted in 10 bins, and then sequenced. Sorted libraries maintained negligible differences in GFP distributions. Weighted average bin numbers according to NGS results were given to each sequence to represent GFP expression. **c**, GFP expression levels of detected sequences with (y-axis) or without (x-axis) Csy4, shown as averaged bin numbers. Shown on the left, q-values, adjusted from t-test’s p-value, were not significant for all sequences. **d**, Inspired anti-CRISPR mechanism. For any processable pre-crRNAs, anti-flanking acrRNAs would repress the processing by forming stacked junctions.

To further prove that long adjacent stems block Csy4 in a sequence-independent manner, we synthesized all 6-bp palindromes without stop codons, or tandem repeat sequences forbidden in DNA synthesis (2689 in total), and 3000 randomly chosen 9-bp palindromes, which were flanked by 4 different sequences (to assure context independence), giving 22756 combinations. These libraries consistently exhibit similar distributions of GFP levels with or without Csy4 (Fig. 3b), suggesting overall protection from cleavage. Indeed, 10-bin FACS-seq (Fig. 3b) showed that GFP expression distributions of pre-crRNAs were highly correlated with or without Csy4 without significant differences as judged by q-values (Fig. 3c). Inspired by this finding of stem-based repression, we opted for designing anti-flanking acrRNAs in order to repress the processing by forming stacked three-way junctions (Fig. 3d).

### Deep learning of pre-crRNA processability for automated design

As a starting point towards repressible CRISPR RNA processing, we set to define the unified rules to identify processable pre-crRNA in absence of cognate acrRNAs. By k-mer analysis, after excluding early-stopped sequences, we found no preferred sequence motifs (Fig. S4a). Thermodynamic^37^ and kinetic variables^40^, and even correct folding of CBS^14^, also showed poor performance as single predictors (Fig. S4b). Therefore, processable pre-crRNAs can be considered as evenly distributed in the sequence space. Given that trivial intra RNA properties failed to depict the rules underpinning this unprecedented sensitivity, it may be that it arises from hitherto uncharacterized 3D structural requirements governing the interactions of the enzyme with its substrate’s flanking regions. Yet, the lack of simple features also raises the difficulty to identify processable pre-crRNAs as potent switches.

To overcome this hurdle, we used supervised learning to build predictive models. The random forest model trained on previously mentioned features failed to predict processable pre-crRNAs though it may accurately identify non-processable pre-crRNAs (Fig. S4d). As a benchmark, we trained 1-dimensional convolutional neural networks (1D CNN) commonly used in previous studies^29, 31^, and used bootstrapping to improve its performance to f1-score ≈ 0.83. To achieve the highest accuracy, we applied a 2-dimensional CNN architecture Seq2DFunc^33^, using a composite matrix representing both sequence and possible base pairings as the input (Fig. 3a), enabling end-to-end prediction. Seq2DFunc significantly outperformed random forest and 1D CNN with f1-score ≈ 0.93 (Fig. 4b). Thus, given acrRNAs with arbitrary sequences, cognate pre-crRNAs can be automatically designed, then approved or rejected by Seq2DFunc, and vice versa (Fig. 4c).

**Fig 4.**
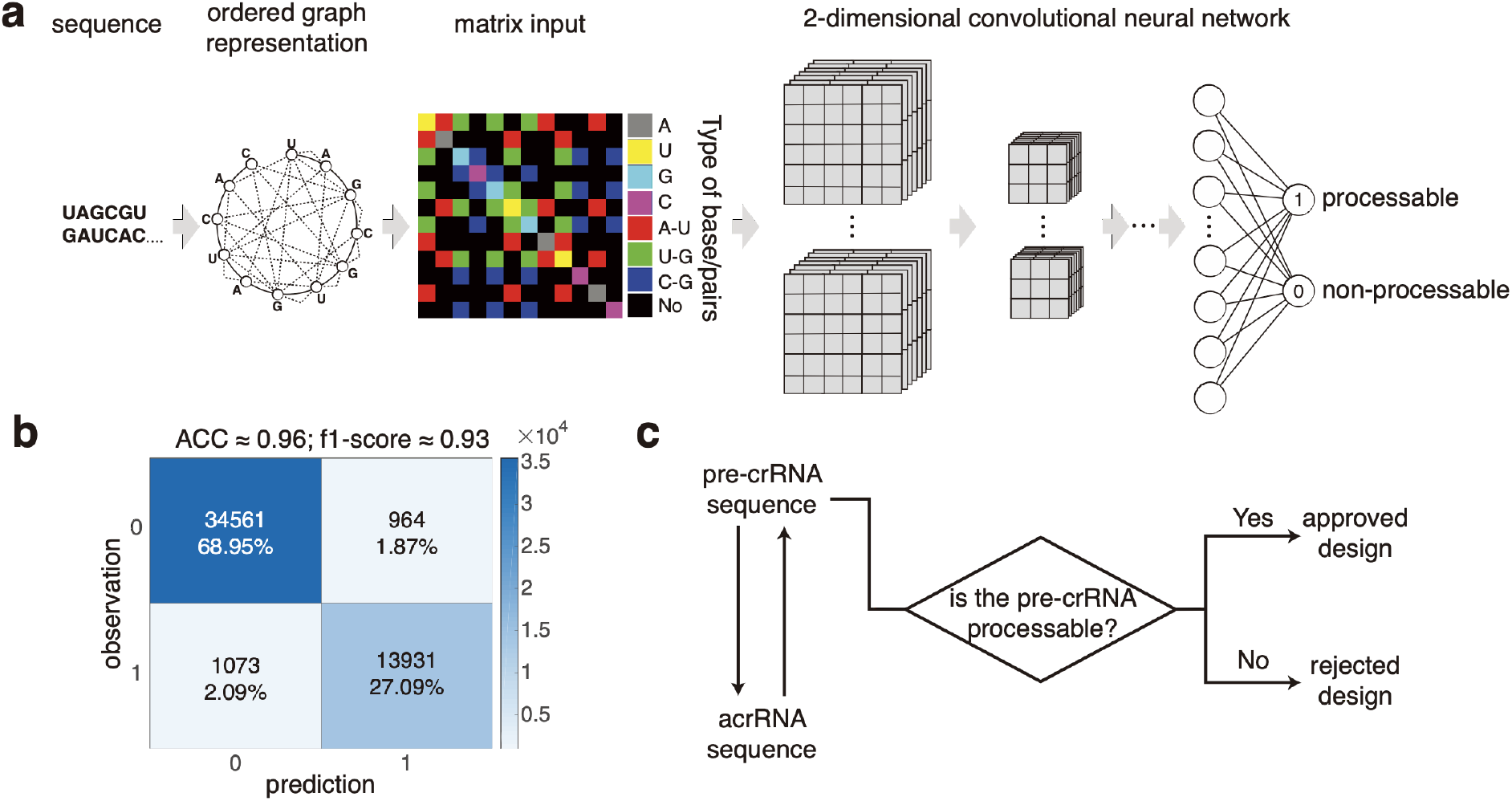
Seq2DFunc architecture to model processability of pre-crRNAs. **a**, Schematic diagram of Seq2DFunc architecture. Sequences were represented as a graph, and then coded by adjacent matrices adding diagonal matrices for sequence information. Matrices were used as inputs to train 2-dimensional convolutional neural networks (see online Methods). **b**, Confusion matrix of the Seq2DFunc models, showing predicted versus observed processability labels. ACC, accuracy. **c**, Process flow diagram of riboregulator designs. Given an acrRNA (or pre-crRNA), we generated its cognate pre-crRNA (or acrRNA), and then evaluated whether the pre-crRNA is processable.

It came to our attention that VIS4Map published simultaneously in an independent preprint^13^ also applied a 2-dimensional convolutional neural network. However, despite some conceptual similarities, the design of Seq2DFunc and VIS4Map were fundamentally different. VIS4Map was trained to visualize structural attentions without incorporating sequence information rather than for accurate prediction. Thus, unlike Seq2DFunc, VIS4Map showed worse accuracy than 1D CNN.

### Designing acrRNAs and pre-crRNAs with deep learning

As a proof of concept, we created an automated pipeline to design 35 cognate pairs of pre-crRNAs and acrRNAs (Fig. 4c, see online Methods). 28 pairs were derived from random sequences. 6 pre-crRNAs were designed to sense thiM riboswitch or mRNA sequences (see online Methods), to represent untranslated RNAs or open reading frames, respectively. For prototyping, we reduced Csy4 expression so that low doses of acrRNAs can result in full repression. We termed GFP production of pre-crRNAs alone “gene ON state” (Fig. 5a). Except one misclassified non-processable pre-crRNA, production rates (see online Methods) were reduced for 2-fold on average upon Csy4 expression, determining the “gene OFF state” (Fig. 5a). All designed acrRNAs fully or partially rescued GFP production of cognate pre-crRNAs (*i.e.*, “gene rescue state”, Fig. 5a). Rescue efficiencies were defined by the ratio of differences between ON-OFF and rescue-OFF states (Fig. 5a). 33 tested pairs of pre-crRNAs showed functional rescue (Fig. 5a, and Table S4), with a significant fraction (16 pairs) exhibiting high (>90%) efficiency (Fig. 5b), validating our pipeline.

**Fig 5.**
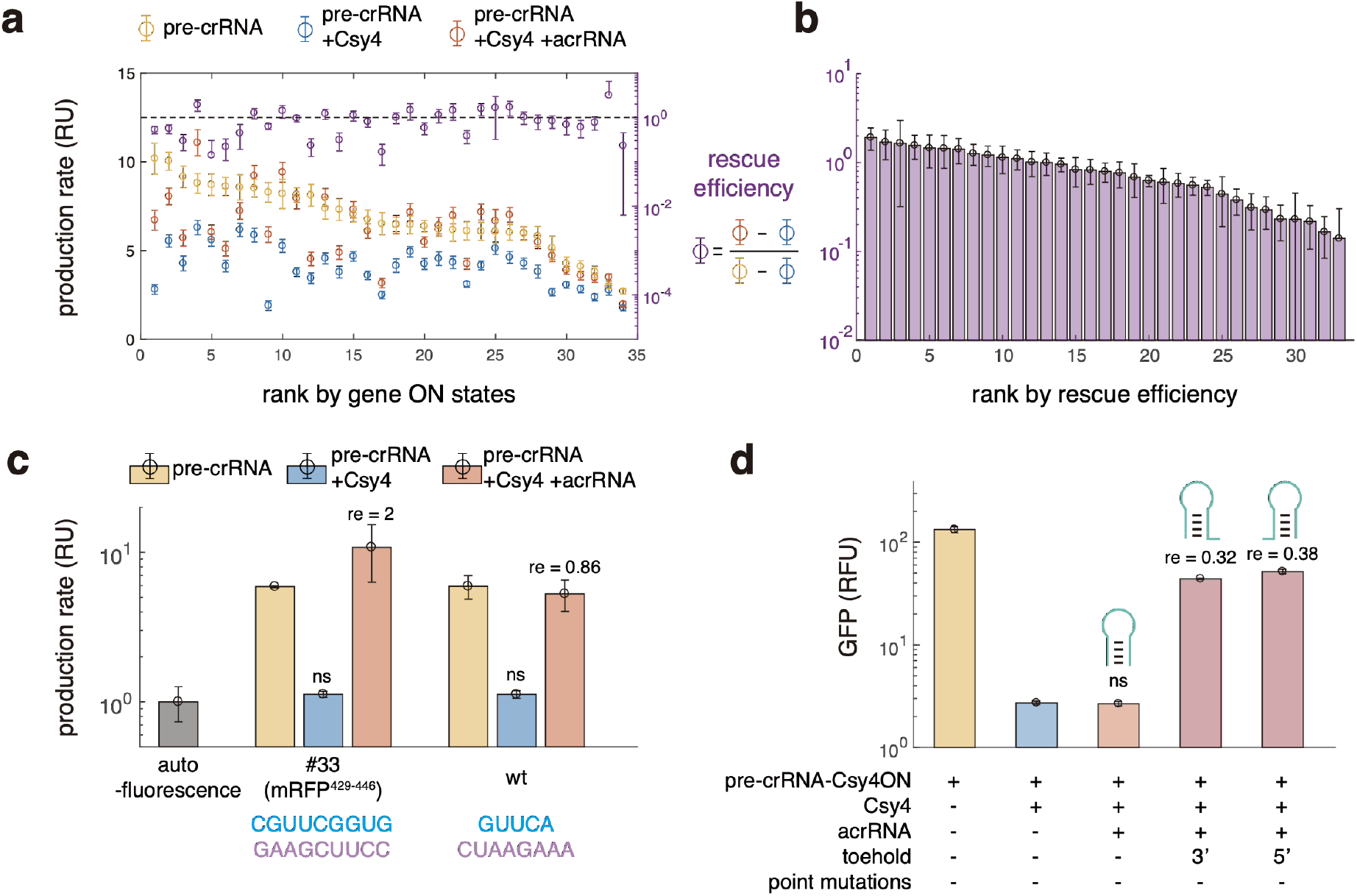
Characterization of anti-CRISPR riboregulatory designs. **a**, GFP production rates of 34 designed pre-crRNAs without Csy4 (yellow), with Csy4 but not cognate acrRNAs (blue), and with both Csy4 and acrRNA (red). RU, relative units, normalized by the autofluorescence accumulation rate. Error bars for production rates, standard deviation of three biological replicates. Error bars for rescue efficiencies were calculated by error propagations. Results were ranked by production rate of pre-crRNA alone, i.e. gene ON states. The rescue efficiency (purple) was defined by the ratio of Csy4-induced GFP expression differences with or without acrRNAs. Design in the dashed box showed no response to Csy4. **b**, Ranked rescue efficiencies of 33 functional pairs of pre-crRNAs and acrRNAs. **c**, Normalized GFP production rates of pre-crRNAs under abundant Csy4 expression without (blue) or with (red) acrRNAs. In both cases, GFP were completely knocked down (blue), and then rescued to full induction levels (yellow) by acrRNAs. ns, the difference between measured sample and autofluorescence is not significant (p-value > 0.05), judged by paired student’s t-test. re, rescue efficiency. **d**, Forward engineering of non-functional acrRNAs of Csy4ON substrate. The acrRNA that only contains the reverse-complementary sequence of Csy4ON’s flanking region displays no function as its stable structure inhibits RNA-RNA interactions. Elongating the anti-flanking regions either at 5’ or 3’ ends rescued the function by providing a toehold domain to initiate RNA-RNA binding. ns, the difference between measured sample and gene OFF state is not significant (p-value > 0.05), judged by paired student’s t-test. re, rescue efficiency.

### Condition optimization for zero-to-one gene activations

With fully rescued pairs, we further increased Csy4 expression to lower gene OFF states for higher-performance gene regulation. With sufficient Csy4, GFP production rates were negligible without leakiness in the absence of acrRNAs, and can be fully rescued by acrRNAs, enabling zero-to-one dynamic ranges (Fig. 5c). Given sufficient expression of acrRNAs for rescue, we anticipate that similar zero-to-one activation can be achieved among designs with different sequences. In comparison, the ranges of ON and OFF states in toehold switches with different sequences vary over thousand-fold and largely overlap^13^, causing large variations in dynamic ranges.

### Forward engineering of acrRNAs for highly structured pre-crRNAs

We found that an acrRNA containing only the 18-nt reverse-complementary sequence of Csy4ON failed to repress the processing (Fig. 5d). It represents a class of insufficient repression due to inefficient RNA-RNA interactions. To overcome this issue, we applied the technique of toehold-mediated strand displacement^41,42^. The original acrRNA were fused with a 15-nt toehold domain that binds to the unstructured sequence in pre-crRNAs. Fusing toehold domain either at the 5’ or 3’ end of the original acrRNA led to significantly higher GFP expressions with similar rescue efficiencies (Fig. 5d).

### Pre-crRNA switches can cover *E. coli* RNAs

Finally, we estimated how programmable the anti-CRISPR riboregulators are. As a demonstration, we used *E. coli* RNAs to act as acrRNAs, and designed pre-crRNAs as switches using machine learning models (see online Methods). We found that about 20% of all possible 18-nt windows in each RNA have a cognate pre-crRNA predicted to be processible. Therefore, on average, each base in *E. coli* transcriptome can be covered 4 times.

## Discussion

Synthetic riboregulators were derived from natural systems using an “antisense” strategy that controls the function of effector domains by direct base pairing^6–8^. Further efforts of toehold switches^9^, toehold repressors^10, 12^, and LASO^10^ opened a window to design riboregulators through an “anti-flanking” mechanism, where effector domains are controlled by context effects. In this study, we further formalized this design paradigm to propose a universal design of riboregulators. For realization, we characterized Csy4-mediated CRISPR RNA processing^14, 22, 23^, and revealed its unusual context-ultrasensitivity, providing binary outcomes in response to flanking structures. We further derived automated designs of riboregulators that repress cognate pre-crRNAs processing by acrRNAs. Lack of design constraints, and consistent performance of this class of riboregulators completed the potential of RNA-based computing. Finally, acrRNA to pre-crRNA signals act on switch pre-crRNAs themselves, enabling conversion of these signals to any downstream events^15–21^. According to profiling of random-flanking libraries and analysis of *E. coli* RNAs, we estimate that about 20-25% sequence in the random space could be available to develop riboregulators. Because non-processable flanking contexts are significantly abundant, other mechanisms to repress RNA processing must exist, which may lead to potential crosstalk issues. Full exploration of rules for repression of crRNA processing therefore would be our interest for future research.

As a result of Csy4’s high affinity and single turnover, anti-CRISPR regulated systems enable dynamic ranges of gene expression from negligible leakiness to full induction. Although we merely illustrated gene activation with large dynamic ranges by controlling the cleavage of coding sequences, we anticipate the great potential of cognate acrRNA and pre-crRNAs coupling as riboregulators in the future, *e.g.,* by gene activation through RBS exposure^21^, or induction of circular RNA formation^20^. Significantly, design of acrRNAs may enable accurate control of type I CRISPR for genome editing^43^ or transcriptional controls^44^. While acr proteins^34, 35^ can only bind to Cas proteins but not distinguish specific crRNAs or guide RNAs, programmable anti-CRISPR RNAs (acrRNAs) may enable crRNA-specific repression, supplementing the anti-CRISPR toolkit. Moreover, the same design principle may be applied to other CRISPR systems such as other Cas6 members or Cas13 that show similar structural requirements or RNA-protein complexes^45^, inspiring more riboregulators.

## Acknowledgements

We are grateful for the support received from C. Lotton and other INSERM U1001 and Center for Research and Interdisciplinarity members, We thank K. Yang (currently in Harvard University) and A. Graham (currently in Imperial College London) for their contributions during the early development of this project, and D. Bikard, Y. Ponty for critical comments on earlier versions of the project. This project has received funding from the European Union’s Horizon 2020 Research and Innovation Programme under the Marie Sklodowska-Curie Grant Agreement No. 665850 and was further supported by the Bettencourt Schueller Foundation.

## Author contributions

H.G conceived the study and experiments, mentored by A.B.L. H.G performed experiments, conventional analysis and random forest models, and conceived the Seq2DFunc. X.S performed the 1D CNN and Seq2DFunc. H.G and X.S analyzed the models. H.G and A.B.L interpreted data and wrote the manuscript with inputs from X. Song.

## Online Methods

### Strain and growth conditions

Unless otherwise noted, DH5a (taxid: 668369) was used for all characterization and routine cloning. MegaX DH10B-T1^R^ (Invitrogen, Carlsbad, CA, USA) was used in large library construction. All strains were grown in LB-Miller medium (BD, Franklin Lakes, NJ, USA) at 37℃ with appropriate antibiotics: ampicillin (100 μg/ml), chloramphenicol (30 μg/ml), and kanamycin (50 μg/ml) for high-copy plasmids and half concentrations for low- or medium-copy plasmid.

### Routine plasmid construction

Plasmid backbone for coding sequence cleavage assay was modified from pRD123 plasmid by Gibson assembly (New England Biolabs). Plasmid backbone for acrRNA cloning was modified from pSB1C3 or BBa_K1100011 from iGEM part registry. To construct plasmids expressing pre-crRNAs and acrRNAs, substrates of interest were assembled by annealing DNA oligonucleotides (Integrated DNA Technologies, Coralville, IA, USA), and then inserted into respective backbones using BsaI-HF®v2 (New England Biolabs) and T4 DNA ligase (Thermo Scientific, Waltham, MA, USA).

### Flow cytometry measurement and analysis

Strains of interest were streaked on LB 2% agar (BD) from glycerol stock and grown overnight at 37℃, and stored at 4℃ after. Single colonies were inoculated into 500 μl LB with proper antibiotics combination, in 2-ml round-bottom 96-well deep-well plates (Thermo Scientific Nunc), and the plate were sealed with polyester acrylate sealing tapes (Thermo Scientific Nunc), and incubated at 37℃ and 750 r.p.m overnight. The overnight cultures were then diluted 250-fold in 500 μl fresh LB medium with antibiotics in 2-ml 96-well U-bottom plate, sealed and incubated at 37℃ and 750 r.p.m for 3 hours to reach log phase. Then the cultures were diluted ~ 900-fold into 100 μl LB medium supplied antibiotics and inducers, in 96-well U-bottom plate (Greiner Bio-One, Kremsmünster, Austria), sealed, and incubated at 37℃ and 750 r.p.m. overnight. Flow cytometry measurements for fluorescence characterization were performed using BD LSRFortessa X-20 cell analyzer with a high-throughput sampler. Cells diluted 10-fold in the phosphate-buffered saline (PBS pH 7.4, Thermo Scientific Gibco) were measured at a rate of 0.5 μl/s. At least 10,000 events gated by forward scatter (FSC) and side scatter (SSC) were recorded for analysis. For each sample, events with negative to zero fluorescence signals (arbitrary units) were excluded, and then medians were used to represent the fluorescence distribution. Mean and standard deviation of median fluorescence were calculated from three biological replicates. Autofluorescence was measured from DH5a strain containing plasmid backbones without fluorescent protein expression. Relative fluorescent unit was calculated by the ratio of fluorescence of each strain over autofluorescence.

### Library design, construction, sorting and next-generation sequencing

Random-flanking pre-crRNA library contained 9-nt random bases on each side of a constant CBS, i.e. N9CTGCCGTATAGGCAGN_9_, were assembled by oligonucleotide annealing. Palindromic-flanking pre-crRNA libraries were designed semi-rationally. All 6-bp and 9-bp palindromes were generated for in-frame insertion. Sequences containing motifs causing synthesis errors were excluded: tandem A_4_, T_4_, G_4_, C_4_, W_6_, S_6_, M_6_, K_6_, R_6_, and Y_6_. Sequences containing stop codons were also excluded. All the rest 6-bp and randomly chosen 3000 9-bp palindromes were used to flank CBS. Resulted sequence were further flanked by 4 types of extra sequences, 5’- or 3’-added TCTTATCTTATCTAT or AGTTTGATTACATTG, to test palindromic-flanking substrates with different contexts that were proven orthogonal in toehold switches^9^. Designed sequences were synthesized as oligonucleotides (Twist Bioscience, USA), and amplified by NEBNext Ultra II Q5 (New England Biolabs). Both libraries were then inserted into the plasmid backbone using BsaI-HF®v2 (New England Biolabs) and T4 DNA ligase (Thermo Scientific). The plasmid libraries were transformed into MegaX DH10B T1^R^ (Invitrogen). A small fraction (~1/20,000) of transformed cells were used to estimate the library size by serial dilution and plating. The rests were diluted 1000-fold, and cultured in LB medium supplied kanamycin and 0.2% arabinose at 37℃ overnight. Overnight cultured cells were diluted into PBS, and then GFP positive cells were enriched using fluorescence activating cell sorting by Bio-Rad S3e Cell Sorter. Sorted cells were cultured overnight and sorted again to minimize the fraction of clones expressing no GFPs. Plasmids extracted from twice-sorted cells were co-transformed with the plasmid expressing Csy4 and mRFP1 into DH5a strain. Transformed cells were similarly cultured overnight and sorted twice by GFP positive or GFP negative gates determined by eyes. Overnight cultures of sorted cells were used for glycerol storages and plasmid extractions. Using forward primer CGTCAAGTTGTCAGGTGG (for random-flanking library) or GCACTACCCGGATGCCTATC (for palindromic-flanking library) and reverse primer GTTCTTCTCCTTTACGACCAG, amplicons were amplified from extracted plasmid library by NEBNext Ultra II Q5 for 15 cycles, and then purified by Wizard® SV Gel and PCR Clean-Up System (Promega, Madison, WI, USA). Purified amplicons were used for sequencing. Next-generation sequencing of the samples was conducted by Plateforme Genomique, Institut Cochin, Paris, France (for random-flanking library) or Genewiz (for palindromic-flanking library).

### NGS sequence mapping and censoring

Identified sequences were mapped to the reference that has the correct length, sequence motif (e.g. CBS, vector sequences). To control the experimental qualities, detected sequences were excluded if reads are lower than the thresholds: for random-flanking library, the total reads in either GFP positive or negative subpopulations should be larger than 3; for palindromic-flanking libraries, the total reads should be larger than 10 both with or without Csy4. Furthermore, for random-flanking libraries, detected sequences were excluded if their GFP positive and negative reads have no significant differences, judged by q-value equal or higher than 0.01. Censored sequences were next analyzed by CD-HIT^46^, to ensure the majority of sequences were at least 2-nt bases away from each other.

### Processability labeling for random-flanking pre-crRNAs

For each identified sequence, the relative frequency of being processed was calculated as the ratio of sum reads in GFP-negative subpopulation (N_(-)_) over the total reads (N_T_), i.e. ρ_(-)_ = N_(-)_ / N_T_. Similarly, the relative frequency of being un-processed was calculated as the ratio of sum reads in GFP-positive subpopulation over the total reads, i.e. ρ_(+)_ = N_(+)_ / N_T_. The processability was defined as arctan2(ρ_(+)_, ρ_(-)_)/arctan2(1,0). Thus, sequences only detected in the cleaved pool have a cleavability as 1, and vice versa. For classification, processabilities lower than 0.3, and higher than 0.7 were further simplified as “0” and “1” respectively, and intermediates were discarded.

### Biophysics and bioinformatics analysis of the RNA substrates

To determine the rules of processing, RNA structural analysis and sequence bioinformatics were applied on the sequences detected from random-flanking libraries. To identify potential sequence motifs, k-mer analysis was applied on the flanking sequences, where k = [1, 2, 3, 4]. Codon usage preferences in *E. coli* (GenScript) were recorded for each flanking sequence. Using ViennaRNA package kinwalker^40^, the folding trajectories of RNA substrates were simulated, and the statistics were recorded including intermediate structures, energy barrier and time required to get into the next structure, and running time. Using Nupack 3.1.0 complex analysis toolbox^37^, RNA structural features under thermodynamic equilibrium were extracted, including the free energy and respective partition function over the ensemble, the number of secondary structures in the ensemble, minimum-free-energy structure (MFE structure) and its energy and equilibrium probability. MFE structures were further characterized by the length of continuous stems, and internal loops as the following. First, whether the Csy4 binding site folded into the 5-bp-5-nt stem-loop was determined. In the range from the upstream flanking context till the fifth base of CBS, the number of bases that pair to the downstream was computed, and vice versa. These two numbers should be the same if CBS folded properly. If CBS was folded in a larger structure, the total number of mismatches was counted as the length of the internal loop. Then the adjacent flanking contexts were analyzed. If the adjacent flanking context forms in extra stems, a length of adjacent stem will be computed from CBS to the first mismatch. If the adjacent flanking context is single-stranded, a length of adjacent loop will be computed from CBS to the first base-pairing. All the computations mentioned above were computed separately for upstream and downstream flanking context, because their structural features are not always symmetric. Individual features were used to fit 70% of the dataset by logistic regression, using Matlab fitclinear function. The resulting models were tested on the rest of the dataset for validation to estimate the prediction power of each feature.

### Random forest model

All the sequence and structural features from the above analyses, including k-mer search, thermodynamic analysis, kinetic analysis, codon usage biases, and mfe structure analysis were integrated into a random forest model with 100 trees, using Matlab TreeBagger toolbox. The random forest model was trained using Matlab Treebagger toolbox with 70% dataset, and then tested on 30% unforeseen testing data.

### 1D CNN model

The nucleotide sequences were one-hot encoded into linear vectors, where A, U, G, C are encoded as [1,0,0,0], [0,1,0,0], [0,0,1,0], and [0,0,0,1]. Vectors were later computed through 1-dimensional convolutions and fully-connected neural networks. The detailed architecture of neural networks is shown in Table S2. 70% dataset were used for training, and then 30% unforeseen testing data were used for testing. Later, we improved 1D CNN by bootstrap sampling the training set. We randomly sampled the entire dataset to construct a training set that had a size as 70% of the total dataset. Sequences not chosen (~50% of database) were used as test sets. Models were coded using Keras package (https://keras.io/), and trained using NVIDIA Tesla P4 GPU.

### Deep learning using Seq2DFunc architecture

For a nucleotide sequence, we consider an ordered graph with N nodes. Each node represents the respective base in the sequence, and is labeled by its content, as A, T (or U), G, and C. Then, all the possible interactions between two nodes are added as edges of the graph. Here, rather than simply applying a complete graph we applied prior knowledge to help the training process. In the following study of RNA elements, we assumed the main pairwise interaction comes from the RNA structure, and thus allowed only Waston-Crick base pairings, A-U, G-C, and a non-canonical base pairing G-U. This choice was guided by the trade off between the efficiency of training, and the potential loss of the information, and depends on researchers’ objective - for instance, higher-order structure such as G-quadruplex will not be represented if only 3 types of base pairs are allowed. Following this, we created a N-by-N matrix, by adding an adjacency matrix indicating all the edges, and a diagonal matrix for all the nodes. To be noticed, the matrix is symmetric. For the label encoding, bases A, U, G, C are encoded as 1, 2, 3, 4, and base pairings A-U or U-A, U-G or G-U, G-C or C-G, as 5, 6, 7. This specific assignment was arbitrarily assigned due to the fact that U and G are capable to pair two types of bases. Commonly-used one-hot encoding was tested but generated too sparse matrices resulting in poor performance, and therefore not shown. 70% of the dataset was used for training, and the rest 30% were used for testing. Bootstrap sampling did not improve the performance. Models were also coded using Keras and trained using NVIDIA Tesla P4 GPU. The architecture of Seq2DFunc is shown in Table S3.

### Automated design pipeline of pre-crRNAs and acrRNAs pairs

For a given RNA sequence as the “seed”, we created a list of all its fragments in an 18-nt window. Then the 18-nt sequences were splitted into two 9-nt pieces to flank CBS. The sequence of flanked CBS (33-nt in total) to construct potential pre-crRNA switches was then evaluated by a valuation model (e.g. Seq2DFunc). Those sequences predicted not to be a processable were excluded. For each remaining pre-crRNA, we designed a cognate acrRNA trigger, which consists of a 5’ stem loop, an 18-nt interaction domain containing anti-flanking sequences, and strong terminator ECK120033737^38^, using Nupack 3.1.0 complexdesign function^37^. We also removed the pre-crRNAs from the list if cognate acrRNAs cannot be designed by Nupack.

### Measurement of GFP production rates

Strains of interest were inoculated and cultured overnight as in flow cytometry measurements. 0.1 μl overnight cultures were replicated into 100 μl fresh medium with antibiotics in μClear 96-well black plate (Greiner, Kremsmünster, Austria), by pin replicators. If Csy4 is driven by inducible PLtetO-1, 25 ng/ml aTc was supplied for low expressions of Csy4. Plates were cultured at 37℃ overnight, and OD600 and GFP were measured in a Tecan SPARK plate reader. GFP production rates were calculated as the partial derivative of GFP with respect to OD600 during the mid-exponential phase of growth by linear regression. Mean and standard deviation of median fluorescence were calculated from three biological replicates. Autofluorescence was measured from DH5a strain containing plasmid backbones without fluorescent protein expression. Relative unit was calculated by the ratio of fluorescence production rate of each strain over the autofluorescence accumulation rate.

---

**Fig. S1.**
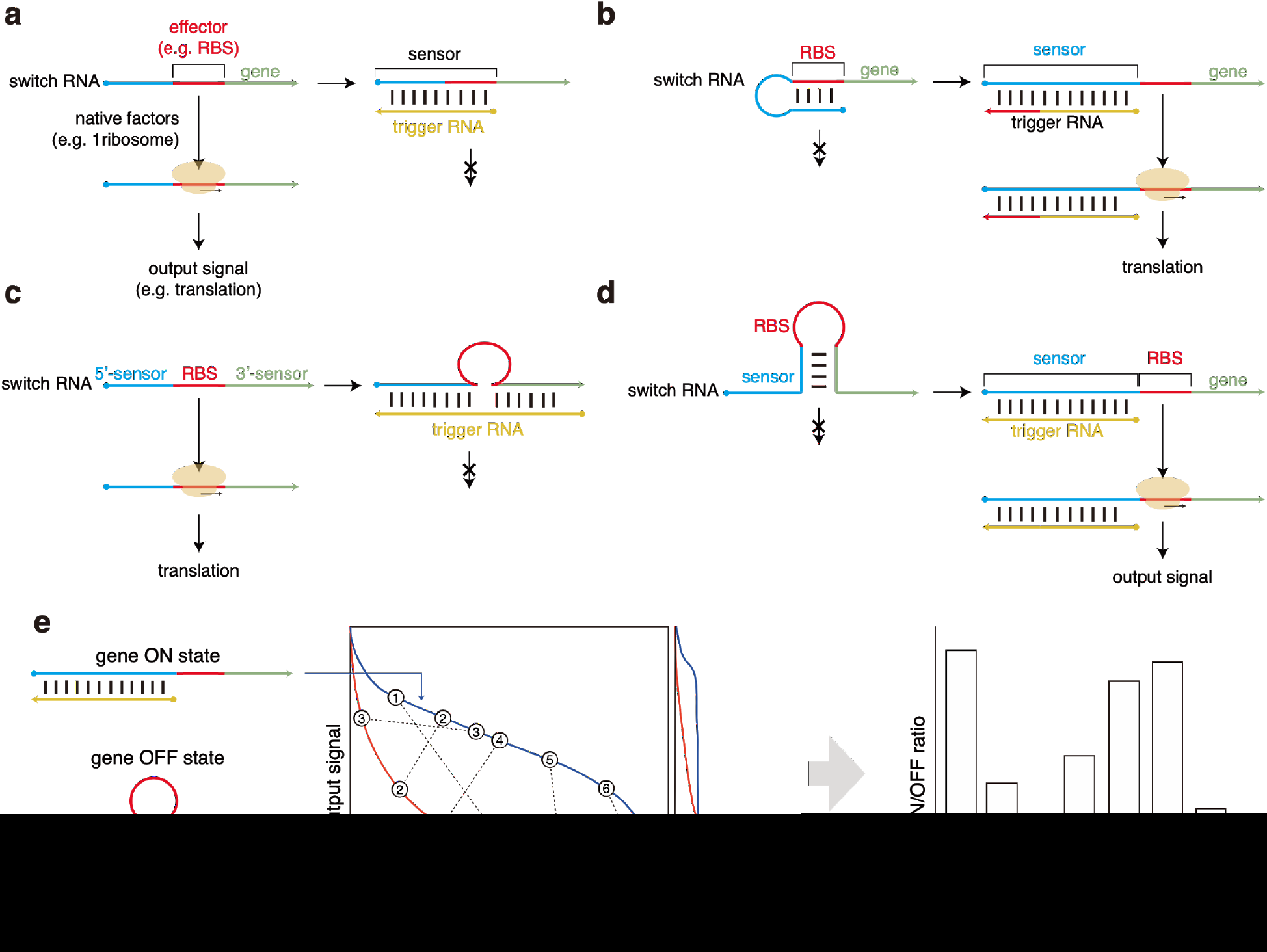
Schematic diagram of previous synthetic riboregulators. **a**, Antisense RNAs^6, 7^. The effector domain, e.g. RBS, drives the function through interactions with native factors (e.g. ribosomes). The switches (i.e. sense RNA) are repressed by triggers (i.e. antisense RNAs) due to base pairing of the effector. **b**, Cis-repressed RNAs^8^. The sensor blocks the effector by base pairing by default, which can be unzipped by cognate triggers. **c**, LASO^10^. compared to antisense RNAs, trigger RNAs (i.e. LASO) now only bind to two-side flanking sequences of RBS and the start codon, and repress their accessibility by context effect. **d**, Toehold switches^9^. a part of the sensor domain base pairs with the 3’-flanking sequence of the RBS except the start codon, to repress the translational initiation by such secondary structure. The rest part of the sensor acts as the toehold to facilitate the trigger binding, which unzips the designed structure and activates the translation. **e**, Schematic depiction explaining significant variations among toehold switches. Conceptually, two designed conformations of toehold switches and switch-trigger complexes should result in gene ON and OFF states. However, their actual outputs may deviate. Using different sequences, the outputs of putative ON and OFF structures span in two open ranges that highly overlap^13^. Therefore, designs with different sequences may present highly varied performance and even be non-functional, shown by hypothetical variants 1-7.

**Fig. S2.**
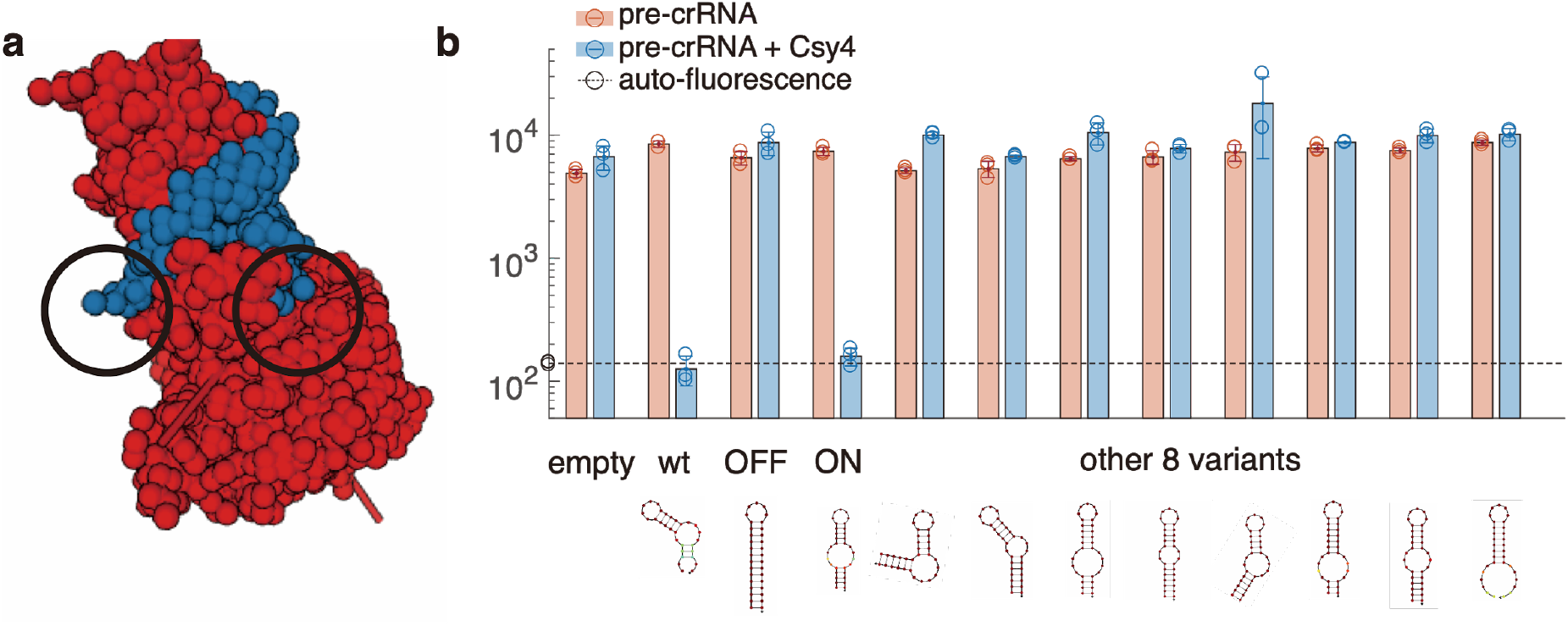
Csy4’s context sensitivity. **a**, the crystal structure of Csy4-CBS complex, PDB 4AL5^22^. Protein and RNA atoms are shown in blue and red. Circles highlight the 5’ and 3’ ends of CBS that are divided by Csy4. **b**, Processing of all rationally-designed substrates. Shown from top to bottom are GFP expression levels without (red) or with (blue) Csy4 under coding sequence cleavage assay, and Nupack-predicted secondary structures of each variant. RFU, relative fluorescence unit, the ratio of arbitrary units between measured sample and autofluorescence. Error bars, standard deviation of three biological replicates.

**Fig S3.**
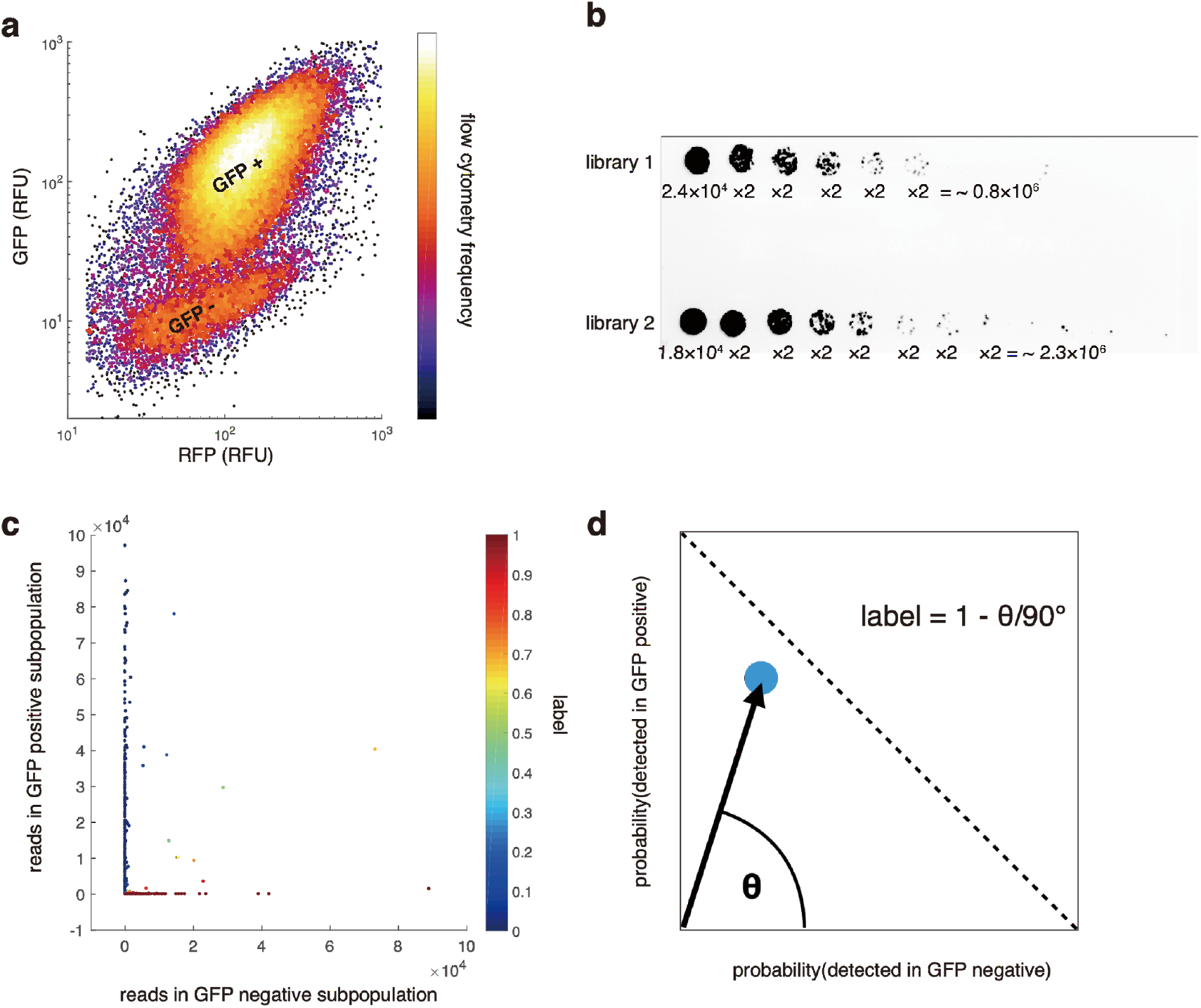
Random-flanking pre-crRNA library analysis. **a**, GFP versus RFP expression levels of cells coexpressing Csy4 plasmids, and pre-crRNA library 2, showing similar two distinct subpopulations. **b**, Estimating library sizes of cells co-expressing Csy4 and indicated pre-crRNA library, by serial dilution and plating. A small fraction of cells co-expressing library 1 and 2 (1 over 24,000 or 18,000, respectively), were serially diluted by 2-folds. Library sizes were therefore estimated as 0.8 and 2.3 million respectively. The agar plate was imaged under RFP fluorescent channel to identify successful transformation of Csy4 plasmids. **c**, Reads detected in GFP negative and positive sorted subpopulations of each detected sequence, colored by processability labels. **d**, Schematic representation of processability. For each sequence, its position in the phase space was plotted according to its probabilities of being detected GFP negative and positive subpopulations. Then, it’s labeled by its angle in the phase space normalized by 0 and 90 degrees, so that sequences only detected in GFP positive were labeled as 0 processable, and vice versa.

**Fig S4.**
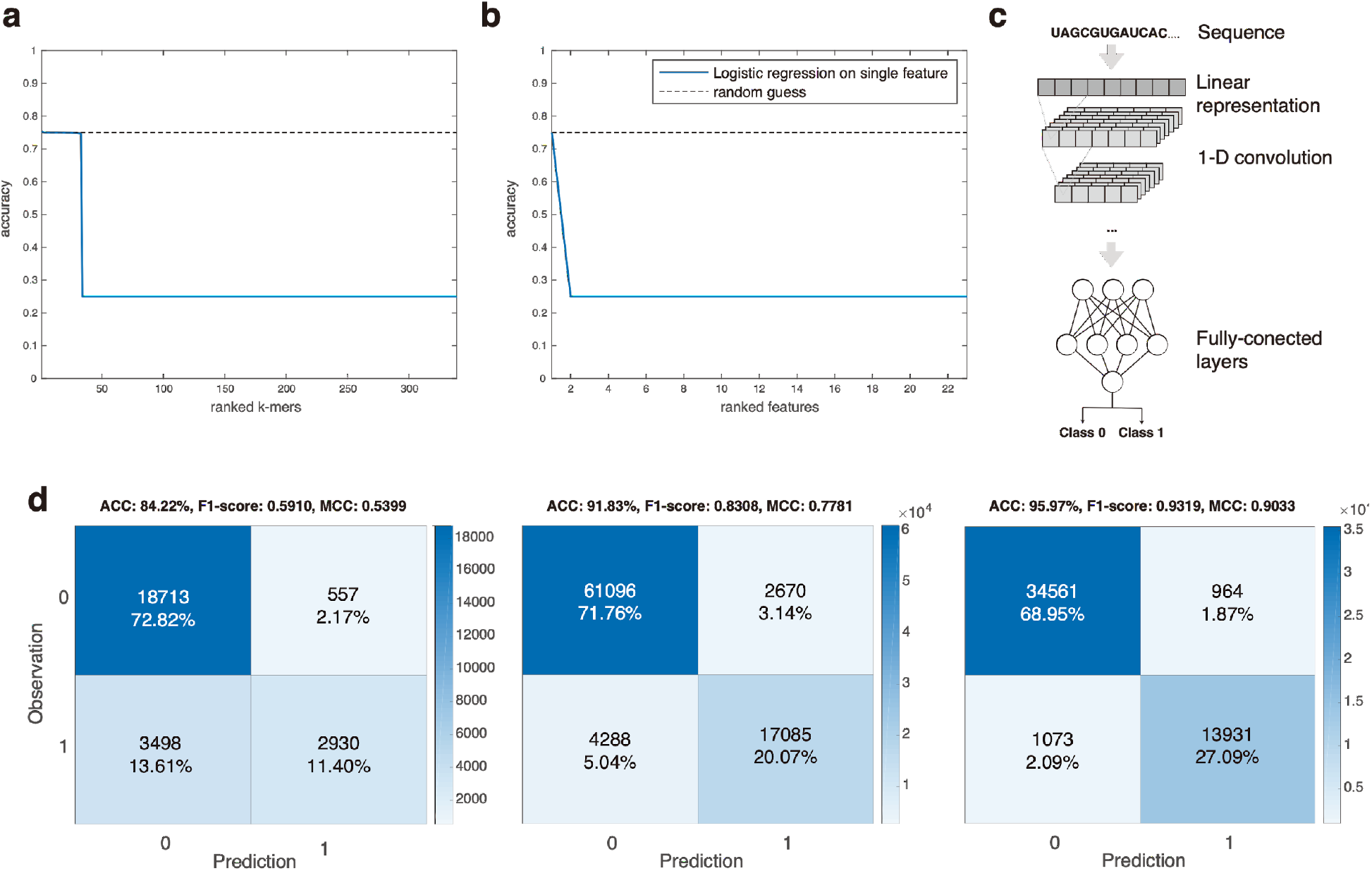
Predicting processable pre-crRNA with feature engineering and deep learning. **a**, Accuracy of logistic regression models using single k-mer information (blue line) compared to random guess (dashed line). **b**, Accuracy of logistic regression models using single thermodynamic and kinetic structural features (blue line) compared to random guess (dashed line) **c**, Schematic representation of 1-dimensional convolutional neural network (1D CNN) that represents the sequence as 1-dimensional vector. **d**, Shown from left to right are confusion matrices of the best random forest, 1D CNN with bootstrapping, and Seq2DFunc models. ACC, accuracy; MCC, Matthews correlation coefficient. Random forest model failed to identify processable pre-crRNAs though it captured the features of most non-processable pre-crRNAs. Both 1D CNN with bootstrapping and Seq2DFunc predicted with high accuracy on unforeseen testing set, and Seq2DFunc outperformed 1D CNN.

**Table S1.**
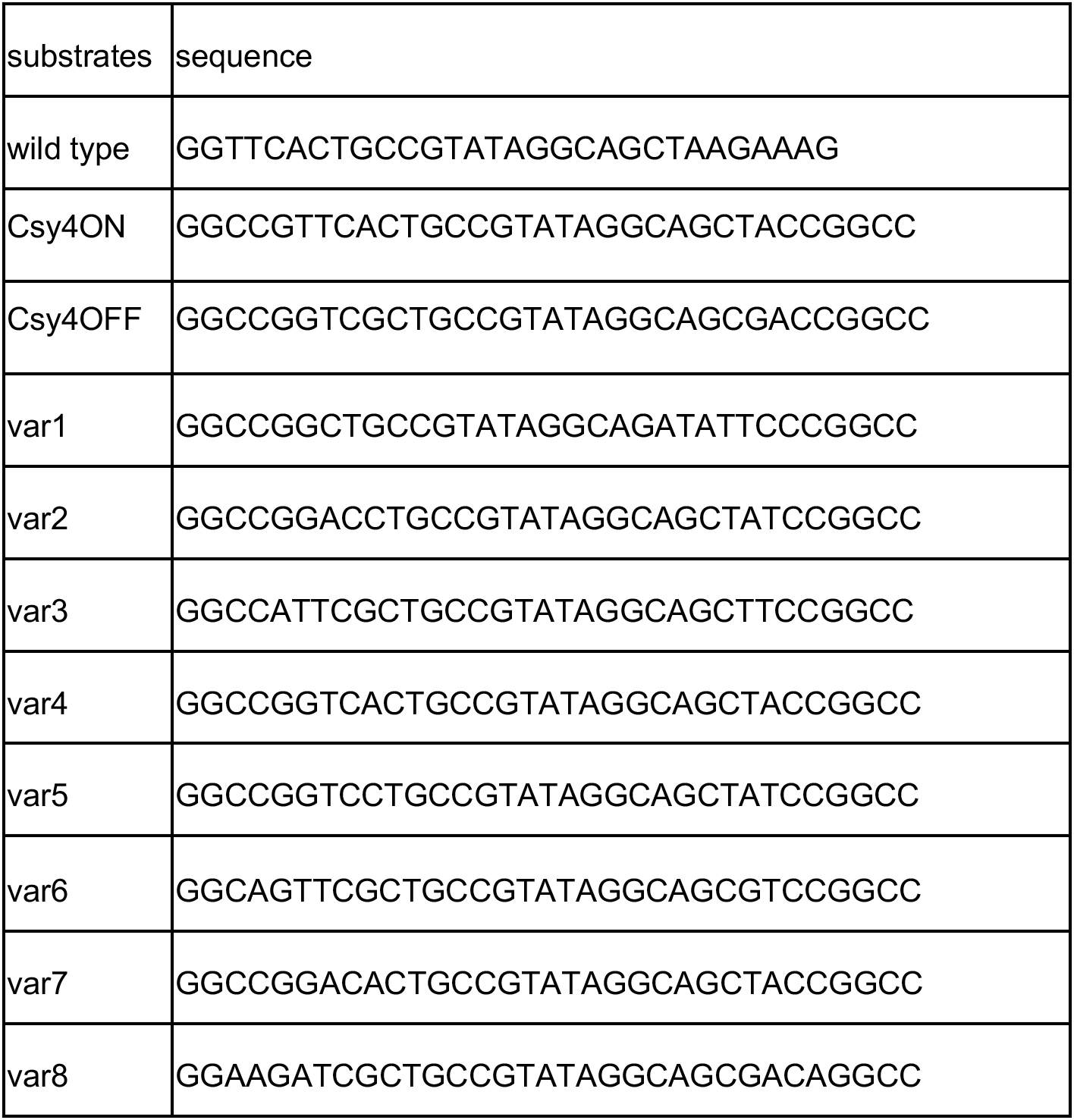
Sequences of rationally designed substrates.

**Table S2.** Architecture of 1D CNN.

| Layer (type) | Output Shape | Param # |
| --- | --- | --- |
| ===== |  |  |
| embedding_1 (Embedding) | (None, 31, 8) | 512 |
| conv1d_1 (Conv1D) | (None, 29, 128) | 3200 |
| conv1d_2 (Conv1D) | (None, 27, 128) | 49280 |
| max_pooling1d_1 (MaxPooling1) | (None, 13, 128) | 0 |
| conv1d_3 (Conv1D) | (None, 11, 256) | 98560 |
| conv1d_4 (Conv1D) | (None, 9, 256) | 196864 |
| max_pooling1d_2 (MaxPooling1) | (None, 4, 256) | 0 |
| conv1d_5 (Conv1D) | (None, 2, 512) | 393728 |
| global_average_pooling1d_1 (GlobalAveragePooling1D) | (None, 512) | 0 |
| dropout_1 (Dropout) | (None, 512) | 0 |
| dense_1 (Dense) | (None, 2) | 1026 |
| ===== |  |  |
| Total params: 743,170 |  |  |
| Trainable params: 743,170 |  |  |
| Non-trainable params: |  |  |
| ===== |  |  |

**Table S3.** Architecture of Seq2DFunc neural networks.

| Layer (type) | Output Shape | Param # |
| --- | --- | --- |
| ===== |  |  |
| conv2d_1 (Conv2D) | (None, 31, 31, 128) | 1280 |
| conv2d_2 (Conv2D) | (None, 29, 29, 128) | 147584 |
| max_pooling2d_1 | (None, 14, 14, 128) | 0 |
| conv2d_3 (Conv2D) | (None, 12, 12, 128) | 147584 |
| conv2d_4 (Conv2D) | (None, 10, 10, 128) | 147584 |
| max_pooling2d_2 (MaxPooling2D) | (None, 5, 5, 128) | 0 |
| conv2d_5 (Conv2D) | (None, 3, 3, 256) | 295168 |
| global_average_pooling2d_1 | (None, 256) | 0 |
| dropout_1 (Dropout) | (None, 256) | 0 |
| dense_1 (Dense) | (None, 2) | 514 |
| ===== |  |  |
| Total params: 739,714 |  |  |
| Trainable params: 739,714 |  |  |
| Non-trainable params: 0 |  |  |

**Table S4.**
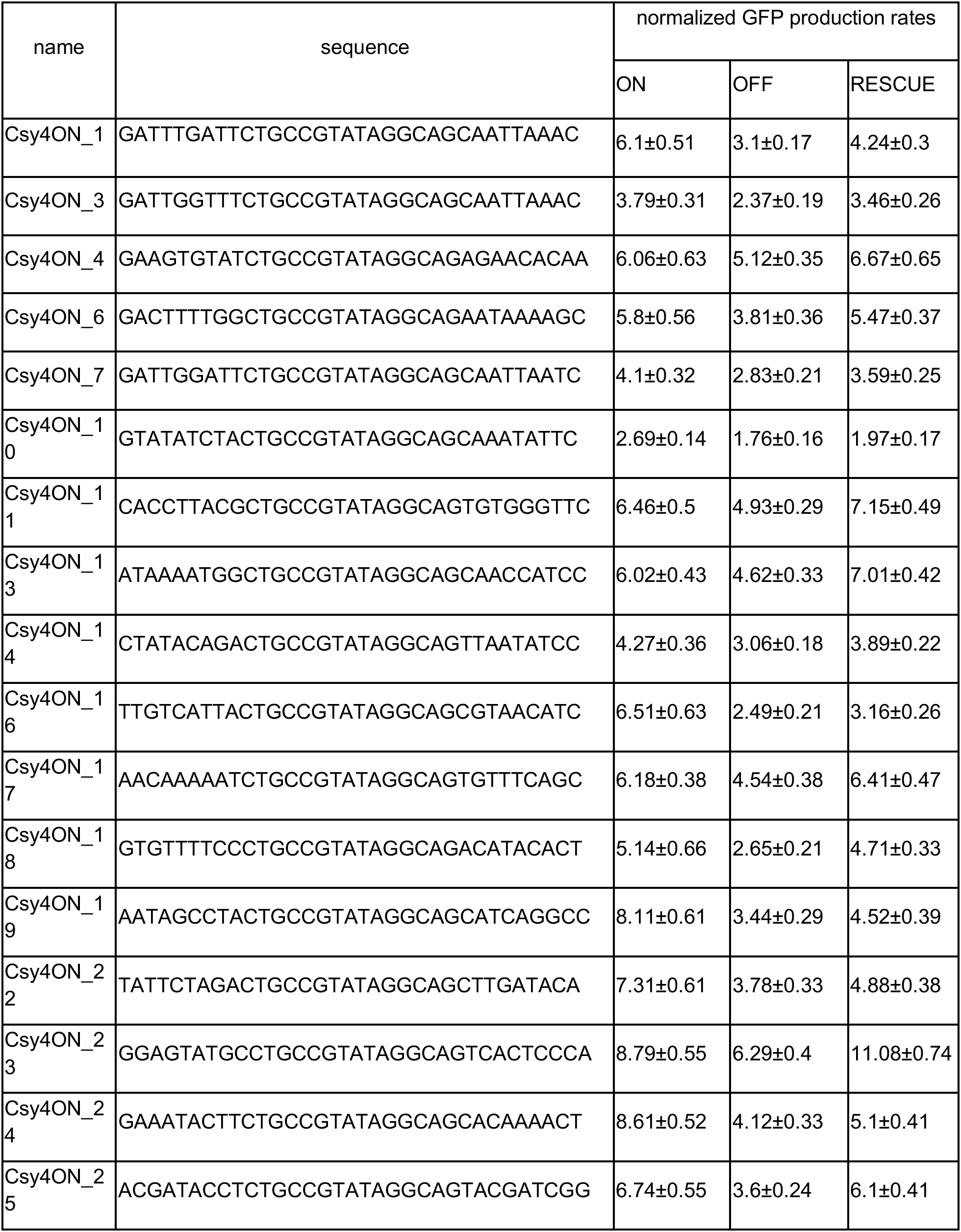
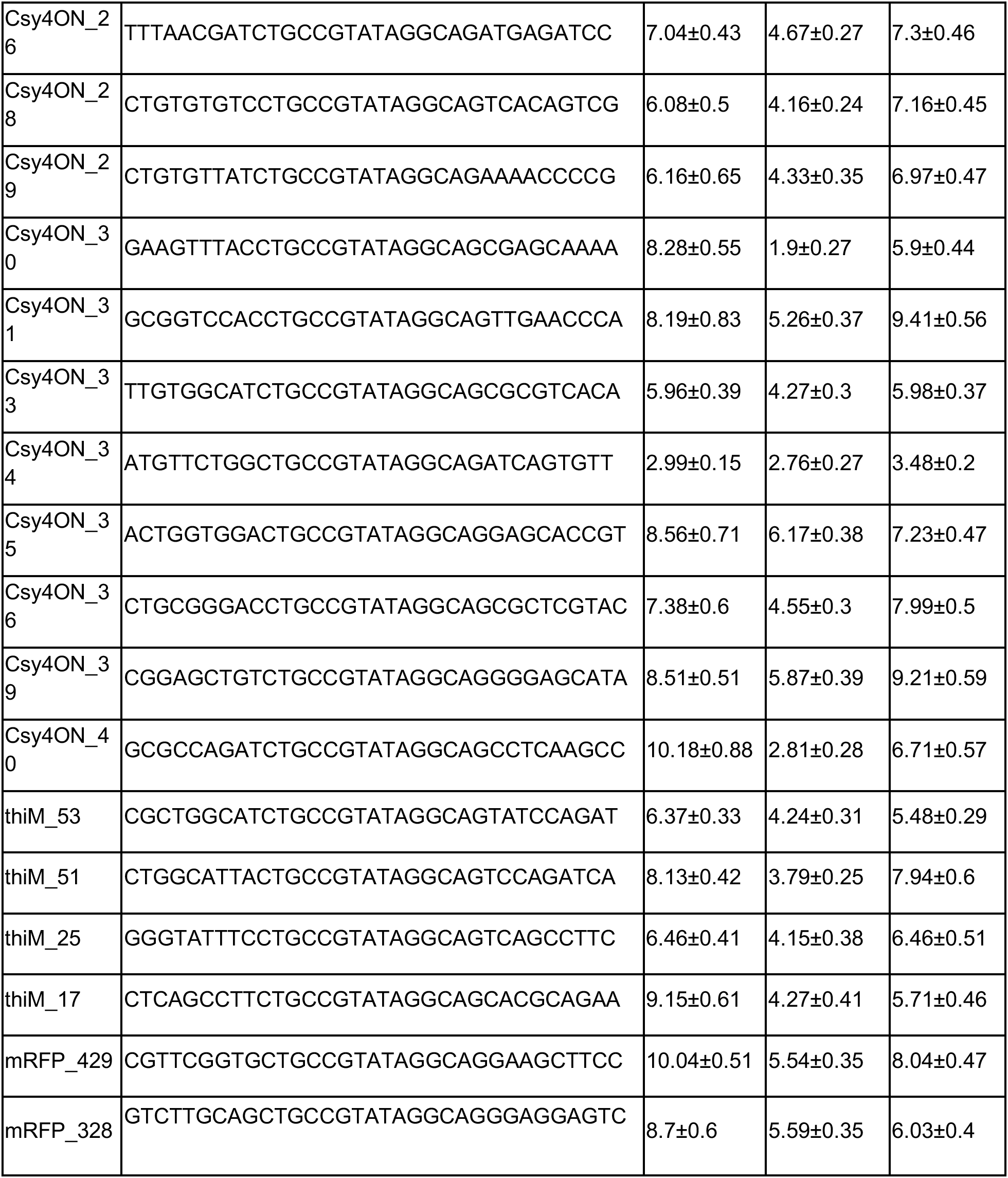
Sequences of pre-crRNAs and respective normalized GFP production rates shown in Figure 4d.

